# Vulnerability and Resilience to Activity-Based Anorexia is Not Sex-Dependent

**DOI:** 10.64898/2026.06.05.730470

**Authors:** Julie Zhao, Jeff A. Beeler, Nesha S. Burghardt

## Abstract

**Introduction:** Anorexia nervosa (AN) is more prevalent in women than men, although rates in men are rising. Animal models can provide insight into whether this differential prevalence is rooted in biological mechanisms, but prior studies have yielded conflicting findings. Using the activity-based anorexia (ABA) model, we previously demonstrated that female mice exhibit distinct vulnerable and resilient phenotypes. Here, we use this phenotypic framework to investigate sex differences in ABA susceptibility.

**Methods:** We tested young adult male C57BL/6N mice using the same ABA protocol used to test age-matched C57BL/6N females. Individual differences in daily bodyweight, food intake, water intake, and wheel running were analyzed and compared across sexes.

**Results:** Males exhibit the same vulnerable and resilient phenotypes as females, with no sex difference in the proportion of mice exhibiting each phenotype or the repertoire of behaviors characterizing them. In both sexes, vulnerable mice exhibit catastrophic weight loss driven by excessive light cycle running, while resilient mice exhibit weight stabilization driven by adaptive changes in consumption. Running during the feeding window revealed that vulnerability is not driven by a decision to run instead of eat in either sex.

**Conclusions:** ABA models adaptive and maladaptive responses to food restriction in both sexes. Behavioral responses to starvation are similar across sexes, suggesting that sex differences in AN prevalence may be driven by stronger sociocultural pressures faced by women to lose weight.

## INTRODUCTION

Anorexia nervosa (AN) is an eating disorder that has the highest mortality rate of any psychiatric disorder (1, 2). It is characterized by an intense fear of gaining weight, a distorted body image, and self-starvation, leading to life-threatening weight loss. Although not part of the formal diagnostic criteria, individuals with AN commonly engage in excessive exercise, which has been implicated in both the development and maintenance of the disorder (3). High levels of premorbid activity increase the risk of developing AN (4) and prolong treatment duration (5, 6), while excessive exercise after discharge is associated with faster relapse (7, 8). Women are reported to be more susceptible to AN than men, with lifetime prevalence rates as high as 4% in women compared to 0.3% in men (9), although rates in men are increasing (10). This disparity may reflect sex differences in the physiological mechanisms underlying AN and/or different sociocultural pressures faced by men and women to be thin (11, 12).

Activity-based anorexia (ABA) is a widely used rodent model of AN that combines restricted access to food with unlimited access to a running wheel. Studies dating back to the 1950s have shown that rats tested under these conditions exhibit a paradoxical increase in wheel running and decrease in voluntary food intake, leading to extreme weight loss and death (13, 14). Investigators have long questioned whether there are sex differences in ABA. It was initially expected that females would be more vulnerable than males (15), because they run more at baseline (16) and higher baseline running had been associated with greater ABA vulnerability (16). In support of this, there are several studies reporting that vulnerability is higher in females (15, 17-19), but others report that females are actually *less* vulnerable than males (20, 21) or that there is no difference between the sexes (22-24). These discrepant findings may be attributable to numerous methodological differences across studies, including variations in the feeding schedule used, number of days of food restriction, and the rodent species and strain used. Perhaps most importantly, studies differed in how ABA vulnerability was determined. While some studies used weight loss and wheel running (17-20, 22, 23, 25, 26), others used survival time (15, 24), with the latter depending on complex physiological changes underlying mortality. Furthermore, few studies identified which behaviors drive weight loss and whether this differs between the sexes.

We previously tested ABA in young adult female C57Bl/6 mice and identified vulnerable and resilient phenotypes, each of which was characterized by distinct behavioral responses and changes in body weight (27-29). While vulnerable mice exhibited the expected failure to eat enough to compensate for energy expenditure, we found that it was their dramatic increase in running during the light cycle that drove their catastrophic weight loss. In contrast, resilient mice initially lost weight, but then adapted by progressively increasing consumption when food was available and decreasing running during the dark cycle, both of which led to weight stabilization. These findings provide a new framework for evaluating susceptibility to ABA, which not only takes into account behaviors driving changes in body weight, but captures individual differences in these responses. With our analysis of phenotypes, we now have the opportunity to compare the responses of males and females in a novel way, potentially providing new insights into sex differences in ABA vulnerability.

Here, we tested ABA in young adult male C57Bl/6 mice and compared their responses to those found in females of the same age, sex, and strain. After analyzing individual differences in food intake, water intake, and wheel running, we discovered that males exhibit the same vulnerable and resilient phenotypes as females. Notably, there was no difference between sexes in the proportion of mice exhibiting each phenotype. Analysis of wheel running during the period of food availability revealed that vulnerability is not driven by a decision to run instead of eat in either sex. Together, our findings indicate that there is no sex difference in vulnerability to ABA, suggesting that the higher prevalence of AN found in women may be due to sociocultural factors rather than biological mechanisms.

## MATERIALS AND METHODS

### Animals

Young adult C57Bl/6N mice of both sexes were purchased at 8 weeks of age (Taconic Biosciences, Germantown, NY) and group housed (4/cage) upon arrival. They were maintained on a 12-hour light-dark schedule with free access to standard laboratory chow (Prolab Isopro 3000 5P75, WF Fisher & Son Inc., Somerville, NY) and water in a temperature-controlled room dedicated to ABA testing. Experiments were conducted in accordance with NIH guidelines and were approved by the Institutional Animal Care and Use Committee of Hunter College.

### ABA Procedure

ABA was tested as previously described (27, 29) with each sex tested separately. One week after arrival, mice were individually housed with unlimited access to food, water, and a wireless running wheel (ENV-044, Med Associates, Inc., St. Albans, VT) that transmitted wheel rotations continuously to a computer. Mice were undisturbed the next day so they could acclimate to their new housing conditions. On the following 3 days, baseline bodyweight, food intake, water intake, and wheel running were recorded (baseline days 1-3). Food was removed from all mice on baseline day 3, two hours after the onset of the dark cycle. On the following 10 days (ABA days 1-10), mice were weighed and water intake was recorded immediately before the onset of the dark cycle. Once the lights turned off, pre-weighed food pellets were immediately placed in the overhead food bin and mice were given 2 hours of unlimited access to food. Mice were 68 days old on ABA day 1. If mice lost at least 25% of their baseline bodyweight (determined on baseline day 3), they were removed from the experiment and characterized as “vulnerable.” Those that did not require removal were characterized as “resilient.”

### Statistical Analyses

Data were analyzed with two-way ANOVA and Bonferroni’s multiple comparisons test using GraphPad prism software (GraphPad, San Diego, CA). A repeated measures ANOVA (RM ANOVA) was used when the same mice were compared at 2 time-points (baseline vs. ABA). Survival curves were analyzed using the log-rank (Mantel-Cox) test. For analyses across days of food restriction, mice were removed at different time point resulting in data sets with missing values. These data sets were analyzed with the linear mixed-effects (LME) model. Significance was accepted for p<0.05. Weight loss data from the female mice used in this study have been published previously (27), but all male data and all sex comparisons reported here are new.

## RESULTS

### Males are as vulnerable to activity-based anorexia as females

At baseline, males (23.35 ± 0.36g) weighed significantly more than females (18.78 ± 0.30g) (t_(36)_ = 9.68, p < 0.0001). To evaluate whether there are sex differences in weight loss during ABA, we compared the bodyweights of males and females (calculated as percent baseline bodyweight) across days of food restriction. We found that both sexes lost substantial weight during the first four days of restriction, but then exhibited weight stabilization and a modest increase in weight by day 10. Analysis of all 10 days indicated no sex difference in this response (Figure 1A, LME model, sex x day, F_(1,32)_ = 0.50, p=0.48; sex, F_(1,36)_ = 1.78, p = 0.19). In both sexes, we found that a subset of vulnerable animals required removal from the experiment, while a subset of resilient mice remained the entire 10 days (Figure 1B). Interestingly, there were no sex differences in survival (hazard ratio, 1.26; 95% CI, 0.54-3.18; p = 0.54) or the proportion of mice exhibiting each phenotype (Figure 1C, χ^2^_(1)_ = 0.99, p = 0.32), indicating that ABA risk is similar in both sexes. Like females (27), we found that vulnerability in males was marked by a progressive, persistent loss of bodyweight leading to removal from the experiment, while resilience was characterized by initial weight loss followed by weight stabilization or even weight gain (Figure 1D). These dynamic changes in bodyweight were further evaluated by analyzing daily change in bodyweight in both sexes. Regardless of sex, vulnerable and resilient mice initially lost weight at a similar rate, but resilient mice gradually stopped losing weight by the fourth day of food restriction, while weight loss was constant in the vulnerable group (Figure 1E, LME model, phenotype x day, F_(1,89)_ = 0.59, p = 0.44; phenotype, F_(1,39)_ = 10.45, p = 0.002; sex, F_(1,39)_ = 1.01, p = 0.32). Interestingly, the rate of weight loss began to differ between the two phenotypes as early as the third day of food restriction. A two-way ANOVA on average daily weight loss confirmed that there were no sex differences within each phenotype (Figure 1F, sex x phenotype, F_(1,34)_ = 3.45, p = 0.07; phenotype, F_(1,34)_ = 109.9, p<0.0001). Together, these results indicate that males exhibit the same two phenotypes that we previously reported in females and that there is no sex difference in ABA vulnerability or resilience.

**Figure 1.**
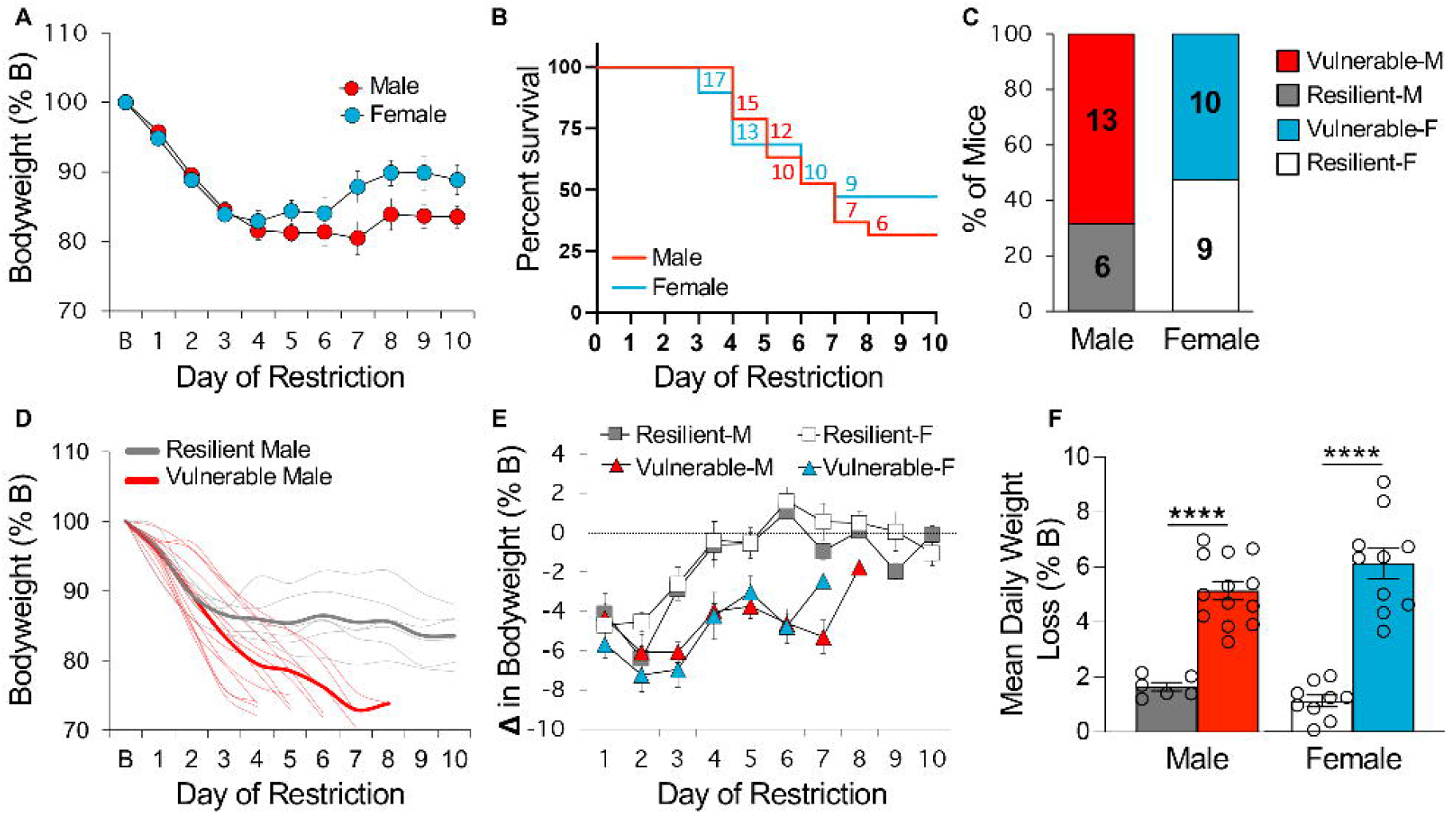
There is no sex difference in ABA vulnerability or resilience. **(A)** Body weight across days of food restriction. **(B)** Survival curves for each sex. Numbers indicate the number of surviving mice that day. **(C)** Percentage of male and female mice that are vulnerable or resilient. Number of animals is indicated inside the bars. **(D)** Body weight of individual resilient (light gray traces) and vulnerable (light red traces) male mice across days of food restriction. Group averages are shown in bold. **(E)** Daily change in bodyweight during food restriction. **(F)** Mean daily weight loss of each group. n = 6 (resilient males, grey), n = 13 (vulnerable males, red), n = 9 (resilient females, white), n = 10 (vulnerable females, blue). Data are represented as mean ± SEM. ****p<0.0001.

### Resilient males adapt food intake

We previously showed that ABA resilience in females is characterized by a progressive increase in food intake (27). Here, we evaluated whether the same is true of males. After comparing vulnerable and resilient males in baseline consumption and consumption during ABA (Figure 2A, timepoint x phenotype, F_(1,17)_ = 24.69, p<0.001), we found that resilient mice did indeed eat more than vulnerable mice during ABA testing (ABA-R vs. ABA-V, Bonferroni, p = 0.0005). Interestingly, there was a trend for resilient males to eat less than vulnerable males at baseline, but this did not reach significance (Baseline-R vs. Baseline-V, Bonferroni, p = 0.058). Analysis of food intake across days of food restriction revealed the resilient males exhibited a steady increase in consumption over time, an adaptive response that was not found in vulnerable males (Figure 2B, LME model, phenotype x day, F_(1,14)_ = 4.23, p = 0.058). Consistent with our previous reports in females (27), only vulnerable males ate less on the day of removal than the previous days of food restriction (Figure 2C, time point x phenotype, F_(1,17)_ = 35.57, p<0.0001, vulnerable before removal vs. vulnerable at removal: Bonferroni, p<0.0001). Finally, changes in bodyweight correlated with consumption in resilient males only (Figure 2D, vulnerable: Pearson r=0.006, p = 0.96; resilient: Pearson r=0.63, p<0.0001). Collectively, these results mirror our previous findings in females and indicate that resilience in both sexes is marked by the same adaptive changes in food intake.

**Figure 2.**
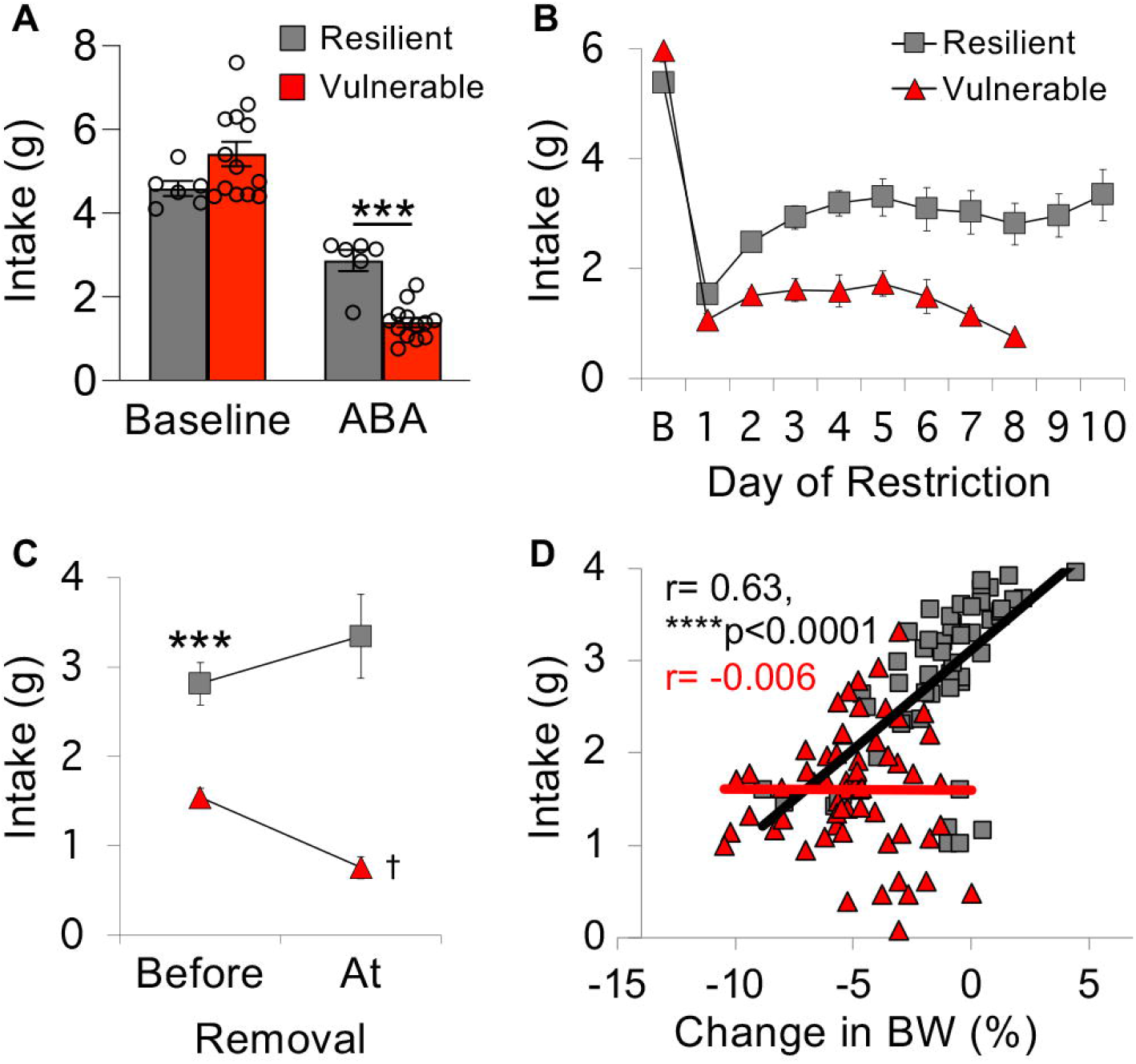
Resilient males exhibit adaptive increases in food intake. **(A)** Average food intake of resilient and vulnerable males before (Baseline) and during 10 days of food restriction (ABA). **(B)** Food intake of resilient and vulnerable male at baseline (B) and on each day of food restriction. **(C)** Average food intake before removal from the experiment and on the last day of testing. ***p<0.001 vs. vulnerable before removal, ^†^p<0.0001 vs. vulnerable before removal. **(D)** Correlation between consumption during restricted access to food and changes in bodyweight (BW) the next day. Each symbol represents 1 animal on 1 experimental day. n = 6 (resilient males), n = 13 (vulnerable males). Data are represented as mean ± SEM. ***p<0.001, ****p<0.0001.

### Weight loss is not driven by reduced water intake in either sex

Food intake often correlates with water intake (30), which also affects bodyweight (31). Here we investigated the contribution of water intake to changes in bodyweight during ABA. In both males and females, water intake did not differ between phenotypes at baseline or during ABA (Supplementary Figure 1A, males, phenotype x timepoint, F_(1,17)_ = 8.06, p = 0.01; vulnerable/baseline males vs. resilient/baseline males: Bonferroni, p>0.05; vulnerable/ABA males vs. resilient/ABA males, Bonferroni, p>0.05; Supplementary Figure 1B, females, phenotype x timepoint, F_(1,17)_ = 2.42, p = 0.14; phenotype, F_(1,17)_ = 3.48, p = 0.08). Closer examination of ABA testing revealed that males of both phenotypes drank less on the day of removal than previous days of ABA testing (Supplementary Figure 1C, phenotype x timepoint, F_(1,17)_ = 0.50, p = 0.49; timepoint, F_(1,17)_ = 5.25, p = 0.035). In contrast, in females, only the vulnerable phenotype exhibited this response (Supplementary Figure 1D, phenotype x timepoint, F_(1,17)_ = 12.73, p = 0.0024; vulnerable/before vs. vulnerable/at removal: Bonferroni, p = 0.02). However, water intake did not correlate with change in bodyweight in any of the groups tested (Supplementary Figure 1E, vulnerable male: Pearson r= 0.14, p = 0.28, resilient male: Pearson r=0.19, p = 0.18; Supplementary Figure 1F, vulnerable female: Pearson r= -0.22, p = 0.19, resilient female: Pearson r=0.06, p = 0.61), indicating that weight loss during ABA is not caused by a reduction in water consumption.

### Vulnerable males exhibit maladaptive light cycle running

We next investigated whether ABA vulnerability in males is associated with the same maladaptive changes in wheel running found in females (27). At baseline, there were no significant differences between phenotypes in amount of running (Figure 3A, LME model: group, F_(1,32)_ = 3.77, p = 0.06) or its circadian distribution (Figure 3B1, LME model: phenotype x hour, F_(1,1347)_ = 1.36, p = 0.24). During ABA, food restriction elicited a similar increase in total running in both phenotypes (Figure 3C, timepoint, F_(1,17)_ = 49.17, p<0.0001; timepoint x phenotype, F_(1,17)_ = 3.64, p = 0.07). However, when light cycle running was analyzed separately, vulnerable males, like vulnerable females (27), exhibited a more dramatic increase in running than resilient mice of the same sex (Figure 3B2, LME model: phenotype x hour, F_(1,3147)_ = 9.70, p = 0.0018; Figure 3D, 3H1, timepoint x phenotype, F_(1,17)_ = 5.47, p = 0.03, Bonferroni, ABA/resilient vs. ABA/vulnerable, p = 0.001). In contrast, dark cycle running increased similarly in both phenotypes during ABA testing (Figure 3E, 3H2, timepoint, F_(1,17)_ = 20.52, p = 0.0003; timepoint x phenotype, F_(1,17)_ = 1.12, p = 0.31). The increase in light cycle running was also more abrupt in vulnerable than resilient males, as indicated by a bigger peak change in running within a 24-hour period (Figure 3F, phenotype x cycle, F_(1,34)_ = 5.55, p = 0.02, Bonferroni, light/resilient vs. light/vulnerable, p = 0.001), which preceded removal from the experiment by 1-2 days for most animals tested (Figure 3G). As previously observed in females, light cycle running positively correlated with weight loss in vulnerable males only (Figure 3I, vulnerable male: Pearson r= -0.23, p = 0.048; resilient male: Pearson r=0.15, p = 0.24), while dark cycle running positively correlated with weight loss in both phenotypes (Figure 3J, vulnerable male: Pearson r= -0.46, p<0.0001; resilient male: Pearson r= -0.30, p = 0.02). These results demonstrate that vulnerability to ABA is characterized by the same maladaptive changes in light cycle running in both sexes.

**Figure 3.**
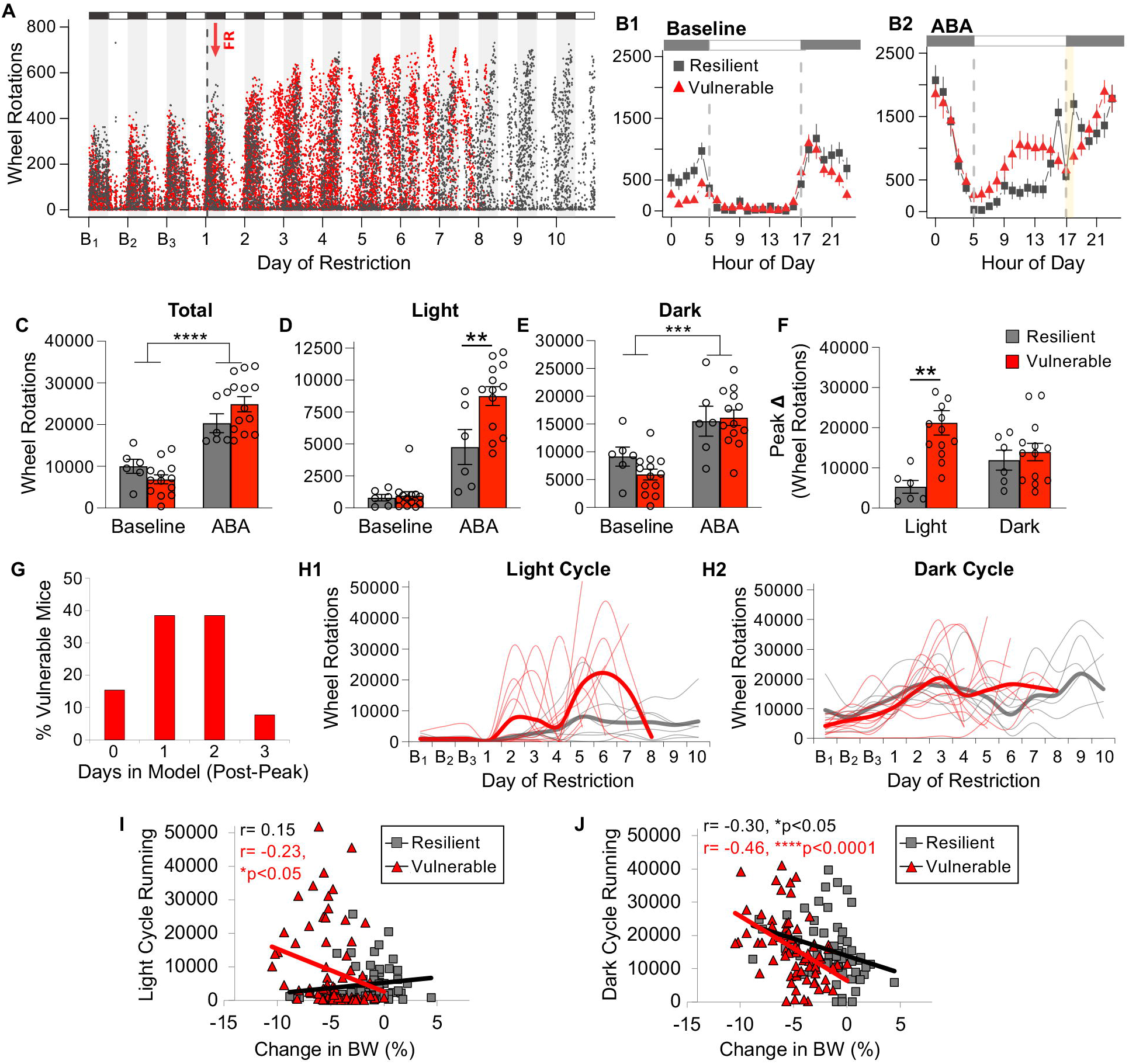
Vulnerable males run excessively during the light cycle. **(A)** Dot plot of wheel running during 3 baseline days (B1-B3) and each day of food restriction for males (resilient, grey; vulnerable, red) in 5-minute bins. Black and gray bars indicate dark and light cycles, respectively. Red arrow indicates start of food restriction (FR). **(B)** Wheel running exhibited by vulnerable and resilient males during **(B1)** baseline days and **(B2)** ABA experimental days in 1-hour bins. Dashed lines indicate the onset (hour 5) and offset (hour 17) of the light cycle. Yellow indicates when food was available. **(C)** Total, **(D)** light cycle, and **(E)** dark cycle wheel running averaged across baseline days and days of food restriction (ABA). **(F)** Maximum increase in wheel running across 2 consecutive days (averaged by group) during the light and dark cycles. **(G)** Histogram showing number of days males remained in the model after exhibiting maximum increase in light cycle running. **(H)** Daily wheel running of individual males (light traces) and group mean (bold traces) during the **(H1)** light cycle and **(H2)** dark cycle (vulnerable, red; resilient, grey). **(I-J)** Correlation between **(I)** light cycle or **(J)** dark cycle running and change in bodyweight (BW) for vulnerable and resilient males. Each symbol represents 1 animal on 1 experimental day. n = 6 (resilient males), n = 13 (vulnerable males). Data are represented as mean ± SEM. ****p<0.0001, ***p<0.001, **p<0.01, *p<0.05.

### ABA vulnerability is not associated with more running during food availability

Previous studies report that animals choose to run instead of eat during ABA (32, 33). To evaluate whether there are differences between phenotypes in this response, we measured running during the two-hour window of food availability. We found that vulnerable females actually run *less* than resilient females and *less* than they did at baseline (Figure 4B, timepoint x phenotype, F_(1,17)_ = 11.19, p = 0.004, Bonferroni, ABA/resilient vs. ABA/vulnerable, p = 0.02, vulnerable/baseline vs. vulnerable/ABA, p<0.0001). Vulnerable males exhibited a similar pattern, although the phenotypes did not differ significantly from each other (Figure 4A, timepoint x phenotype, F_(1,17)_ = 1.88, p = 0.19, Bonferroni, vulnerable/baseline vs. vulnerable/ABA, p = 0.045, ABA/resilient vs. ABA/vulnerable, p = 0.31). This occurred despite females exhibiting 2-3 times more baseline running in a 24-hour period than males (Figure 4C, sex, F_(1,34)_ = 54.33, p<0.0001). The reduction in wheel running during the 2-hour feeding window primarily occurred during the end stages of weight loss. Indeed, vulnerable mice of both sexes ran less than same-sex resilient mice on the day they were removed from the experiment, but not earlier in ABA testing (Figure 4D, timepoint x phenotype, F_(1,17)_ = 11.68, p = 0.003; Bonferroni, at removal/resilient male vs. at removal/vulnerable male, p = 0.0004; Figure 4E, timepoint x phenotype, F_(1,16)_ = 6.12, p = 0.03, Bonferroni, at removal/resilient female vs. at removal/vulnerable female, p = 0.0004). Finally, there was no correlation between running during the feeding window and daily weight loss in vulnerable mice of either sex (Figure 4F, vulnerable male: Pearson r= -0.15, p = 0.25; Figure 4G, vulnerable female: Pearson r= -0.13, p = 0.43). However, in resilient females, weight gain significantly correlated with less running during this time window (Figure 4G, resilient female: Pearson, r= -0.29, p = 0.008), results that are in line with the adaptive decreases in dark cycle running previously described (27). This correlation was not found in resilient males (Figure 4F, resilient males: Pearson, r= -0.09, p = 0.50). Together, these findings reveal that vulnerable mice do not run more than resilient mice when food is available, indicating that ABA vulnerability is not driven by a decision to run instead of eat in either sex.

**Figure 4.**
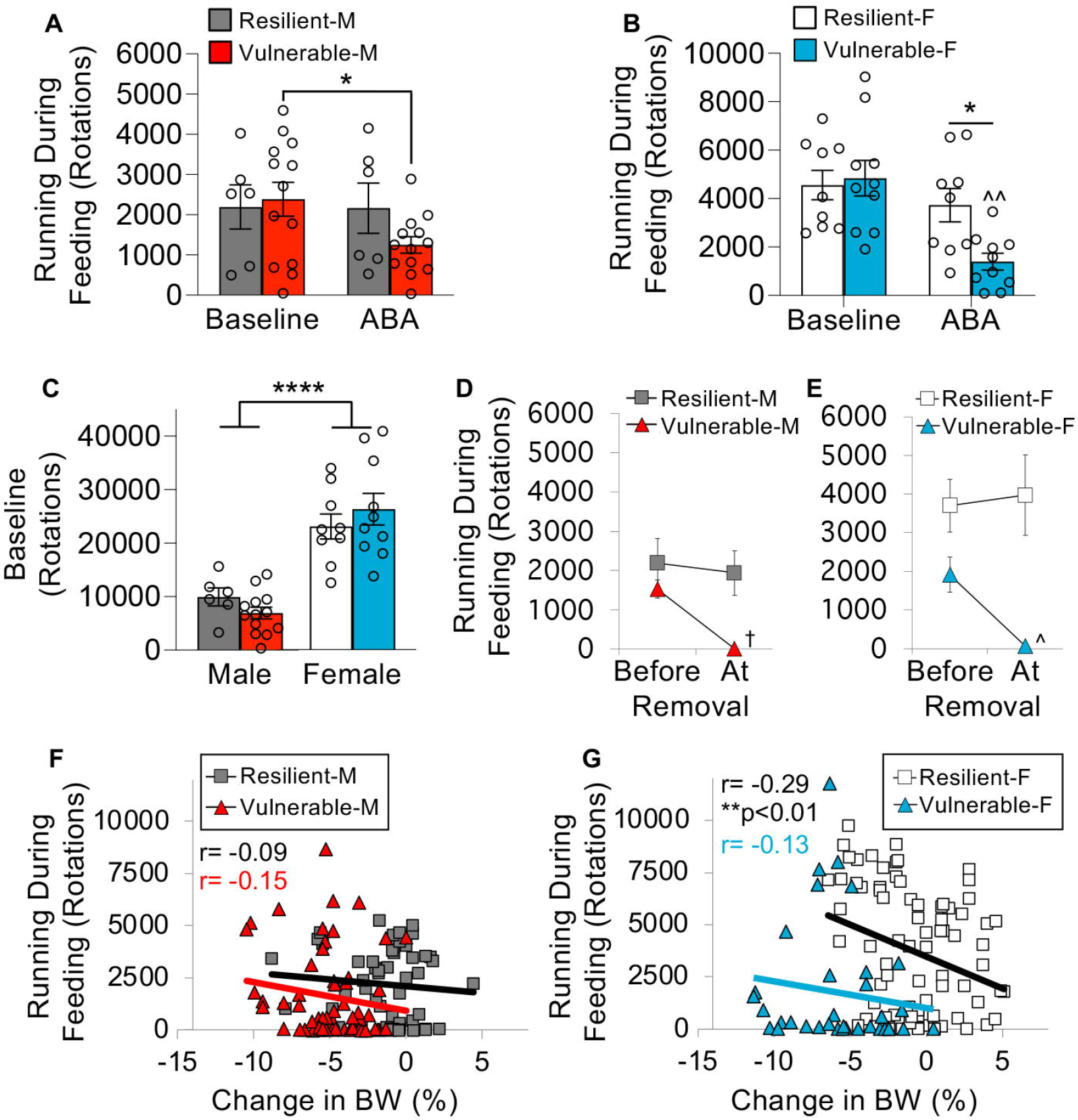
ABA vulnerability is not associated with more running during food availability. **(A-B)** Wheel running exhibited by **(A)** males and **(B)** females during the 2-hour window of food availability (ABA) and during the same 2-hour period before food was removed (Baseline). **(C)** 24-hour wheel running exhibited by males and females averaged across 3 baseline days. (**D-E**) Average running during 2-hour window of food availability exhibited by **(D)** males and **(E)** females before removal from the experiment and on the last day testing. **(F-G)** Correlation between running during 2-hours of food availability and changes in bodyweight the next day for **(F)** males and **(G)** females. Each symbol represents 1 animal on 1 experimental day. n = 6 (resilient males), n = 13 (vulnerable males), n = 9 (resilient females), n = 10 (vulnerable females). Data are represented as mean ± SEM. ****p<0.0001, *p<0.05, ^^p<0.0001 vs. vulnerable female baseline, ^†^p<0.0001 vs. resilient males at removal, ^p<0.0001 vs. resilient females at removal.

## DISCUSSION

We tested ABA in C57Bl/6 male mice and found that they demonstrate the same distinct vulnerable and resilient phenotypes as age-matched C57Bl/6 females. We found no sex difference in the repertoire of behaviors characterizing each phenotype or the proportion of mice exhibiting vulnerability or resilience. Phenotypes were distinguishable based on daily changes in bodyweight that were driven by either excessive light cycle running (vulnerable phenotype) or increased food intake (resilient phenotype), none of which varied by sex. Together, our findings indicate that there are no sex differences in ABA, suggesting that ABA may be used to model adaptive and maladaptive responses of both men and women once they reach the same level of weight loss. The higher prevalence of AN in women may therefore be attributed to stronger sociocultural pressures to lose weight rather than sex-specific biological mechanisms.

Our findings are consistent with ABA studies conducted in Wistar rats reporting that the percentage of baseline bodyweight lost across days of food restriction did not differ between males and females (22, 23)]. In studies that allowed rats to die over the course of ABA testing, there was also no sex difference in survival duration (18, 24) but see (15). Interestingly, one of these studies found that female rats died earlier than males, but concluded that there was no sex effect after beginning bodyweight was included as a covariate in the analysis (18). To control for sex differences in initial bodyweight, another group included older female Wistar rats with weights that were comparable to those of males (23). Older females lost weight more slowly and met removal criterion later than younger mice of both sexes, consistent with the known protective effects of age on ABA vulnerability (27, 34). Like our study in mice, there were no differences between same-age male and female rats, despite males weighing more at baseline (23). Another study used male Sprague-Dawley rats that were only slightly younger than females, in which case the initial weight difference was minimized, but still significant. Contrary to the findings described above, those younger males required removal from the experiment *earlier* than females (20), which again may have reflected an effect of age. Furthermore, that study excluded rats that never required removal (i.e., resilient) from all analyses (20), which may have affected experimental outcome. Our analyses of both vulnerable and resilient mice indicate that starting weight, which was higher in males of both phenotypes, is unrelated to mean daily weight loss or survival in the experiment.

Historically, ABA has been tested primarily in rats, resulting in far fewer mouse studies investigating sex differences. We are aware of only two sex difference studies that used the same mouse strain we used (C57Bl/6), both of which progressively limited food access, which is different from the fixed feeding schedule we used. Interestingly, one study found that C57Bl/6 males lost more weight than females (21), indicating higher ABA vulnerability in males, while the other concluded the opposite, with C57Bl/6 males surviving in the model *longer* than females (19). Notably, both challenged mice in ways that were fundamentally different from our study. While they allowed mice to gradually adapt to a restricted feeding schedule, we evaluated responses to a sudden and consistent change in food access. They also gave mice more time to eat, with food access that was eventually limited to 3 hours compared to the 2-hour feeding window we provided. In addition, they removed mice from the experiment once they lost 20% of their baseline weight for 2-3 consecutive days (19, 21), which was less strict than our removal criterion (> 25%). If we had used their criterion, some of our resilient mice that lost 20% baseline weight would have been removed before they had a chance to show weight stabilization. While these methodological differences may account for our inability to replicate either of the previously reported sex differences in this strain, it remains unclear why those previous findings were the opposite of each other.

Interestingly, when ABA was tested in Balb/cJ mice, which are more anxious than the more commonly used C57Bl/6 strain (35), males were found to survive longer in the model than females (17). Such findings highlight the role of mouse strain in ABA vulnerability (17, 36-38) and implicate a potential sex difference in how anxiety might affect ABA.

Consistent with numerous reports in mice (19, 25) and rats (16, 39), we found that females run significantly more than males before food is restricted (i.e., baseline). These baseline differences can affect the interpretation of sex differences during ABA, where higher running in females could reflect a stronger effect of food restriction and weight loss on activity or a sex difference that always existed. Given that more baseline running has been associated with stronger ABA susceptibility (16, 39), the expectation has been that females would be more vulnerable to ABA than males (16). However, our findings demonstrate that there is no sex difference in ABA vulnerability or resilience and that mice of both phenotypes exhibit similar levels of running at baseline. In one mouse study, the association between baseline running and weight loss was based on 4 days of food restriction (39), which is shorter than our 10-day protocol. It is possible that baseline running predicts early changes in bodyweight that are later corrected in resilient mice, potentially accounting for why baseline running did not differ between phenotypes. Indeed, we show that resilient mice stopped losing weight after the fourth day of food restriction, which is after data collection ended in that study.

Finally, we analyzed running during food availability to evaluate whether ABA vulnerability is characterized by a decision to run instead of eat (40-42). However, we found that, if anything, vulnerable mice run *less* than resilient mice during this time window, findings that are in line with a previous study reporting decreased running during feeding in both sexes (19). We also found no association between running during the feeding window and weight change in vulnerable mice, further indicating that this is not what differentiates vulnerability from resilience. Instead, we found that excessive running throughout the light cycle played a key role in driving weight loss in vulnerable mice, effects evident in both sexes.

Based on known sex differences in the prevalence of AN, the majority of AN studies have been conducted in women only. Similarly, ABA studies frequently use female rodents, with much less is known about the cellular and molecular basis of ABA in males (40). However, the rates of eating disorders in males have been increasing (10) and are higher than previously thought (43). A comparison of AN in men and women reveal that core symptoms, such as patterns of food restriction, are similar in both sexes (44), but food restriction is often initiated for different reasons. While women are focused on being thin, men tend to be more concerned with muscle definition (43, 45, 46) and men engage in more excessive exercise (47). Our ABA findings suggest that once sufficient weight loss occurs, the likelihood of expressing an adaptive (i.e., resilient) or maladaptive (i.e., vulnerable) response is not sexually dimorphic. Sex differences in the prevalence of AN may therefore reflect stronger sociocultural pressures faced by women to be thin, driving more women to reach levels of weight loss that trigger onset of the disorder. Furthermore, men are less likely to seek treatment than women when they experience comparable levels of problematic eating behaviors (47), potentially skewing prevalence data. Future ABA studies identifying substrates underlying vulnerability and resilience may provide important insight into the etiology and maintenance of AN in both sexes. Notably, investigation of factors mediating ABA resilience, defined by maintenance of bodyweight rather than slower decline of bodyweight, may contribute importantly to our understanding of factors protecting individuals who diet and exercise from developing AN. Such information may not only lead to the development of novel treatment options, but the identification of biomarkers for early diagnosis in both sexes.

## Acknowledgements

This work was supported by a Klarman Family Foundation Eating Disorders Grant (JB and NB) and PSC-CUNY Awards jointly funded by the Professional Staff Congress and The City University of New York (JB and NB).

