## Supplementary figures and images for "Vulnerability and Resilience to Activity-Based Anorexia is Not Sex-Dependent"

### Supplementary Figure 1

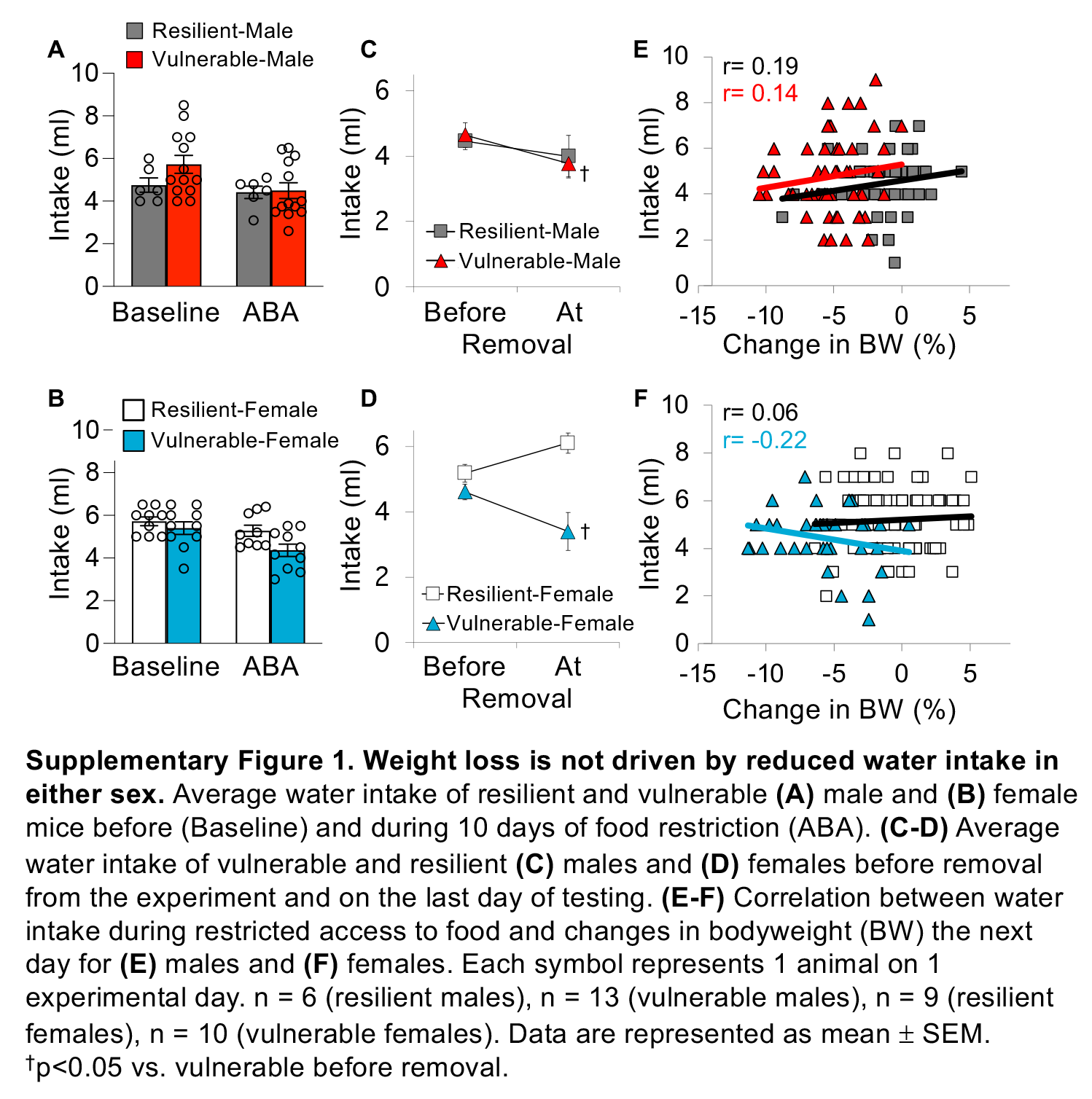
